# Development and Validation of the Transcranial Magnetic Stimulation Reporting Assessment Tool (TMS-RAT)

**DOI:** 10.64898/2026.02.18.706460

**Authors:** Orsolya Székely, Nicholas P. Holmes, Jennifer Ashton, Friederike Breuer, Hsin-Yuan Chen, Nunzia Valentina Di Chiaro, Arnaud Duport, Polytimi Frangou, Louisa Gwynne, Umair Hassan, Cassandra J. Lowe, Brian Mathias, Nan Peng, Jessica L. Pepper, Phivos Phylactou, Malgorzata Anna Szymanska, Luigi Tamè

## Abstract

**Highlights:**

- We introduce the TMS-RAT, a reporting (assessment) tool for TMS studies
- Developed within a community-informed, iterative process rating 333 TMS studies
- Empirically evaluated for usability, inter-rater, and test-retest reliability
- A validated subset enables reliable retrospective assessment of reporting
- The modular structure enables use across a wide range of TMS study designs

**Background:** A standardised tool for comprehensive reporting can improve transparency, support consistent documentation, and enable comparison across transcranial magnetic stimulation (TMS) studies. The most used reporting checklist lacks clear definitions of full reporting and was not initially evaluated for usability or inter-rater reliability. A scoping review of studies using this checklist shows that its items are reported only 50% of the time, suggesting that method descriptions are often incomplete.

**Methods:** We developed the TMS Reporting Assessment Tool (TMS-RAT), a comprehensive reporting framework that provides clear definitions and examples for its items, covering a wide range of TMS protocols. We tested the usability and reliability of the TMS-RAT by rating all studies published between 1991 and 2025 using afferent conditioning (n = 333), a protocol encompassing many reporting categories identified during tool development. Seventeen independent raters contributed across three development phases, a validation phase, and a retest phase, with naïve raters introduced in each phase. Iterative refinements of the tool were informed by inter-rater reliability, qualitative rater feedback, and consultation with external TMS experts.

**Results:** We present two versions of the tool: the 72-item TMS-RAT v1.0, designed to guide comprehensive reporting, and the TMS-RAT v1.1, a subset of 50 items with the highest inter-rater (overall AC1 = 0.78, range = [0.60–0.99]) and test-retest reliability (overall AC1 = 0.82, range = [0.65–1.0]), intended for retrospective evaluation of reporting in systematic reviews, meta-analyses.

**Conclusion:** The TMS-RAT is a comprehensive, reliable tool that seeks to improve transparency and reproducibility in TMS research.

## 1. Introduction

### 1.1 Background

For transcranial magnetic stimulation (TMS) studies to contribute maximally to scientific progress, their core methods have to be reported clearly enough for others to interpret the results accurately, build on the findings, or reproduce experiments across laboratories and over time. For over forty years, TMS has been used to investigate the human nervous system (Barker et al., 1985), but the field has often struggled with inconsistent effects and difficulties replicating earlier findings (Hamada et al., 2013; Héroux et al., 2015). A standardised tool for reporting (and assessing the reporting of) core methods can guide the design and documentation of TMS experiments, and enable retrospective assessment and comparison of reporting completeness in systematic reviews and meta-analyses.

The most commonly used TMS reporting checklist was developed by Chipchase et al. (2012), using a two-round Delphi consensus process (Hsu & Sandford, 2007) during which 42 nominated experts selected and prioritised items considered essential to report or control. This 30-item checklist provided the foundation for reporting and assessing TMS studies, but has several limitations. First, Chipchase’s checklist was designed to assess studies of the motor system exclusively and has limited application to other TMS domains. Second, the checklist lacks definitions or examples of what constitutes full reporting. For example, participant age may be reported as ‘adults’, a range, a group mean (with or without a standard deviation; SD), or as each individual participant’s age. Without full, consistent reporting, it is difficult to evaluate age-related effects or confounds across studies in meta-analyses. Third, the checklist was not initially evaluated for usability or inter-rater reliability. When this was assessed in later reviews, about one third of checklist items did not achieve acceptable inter-rater agreement or precision, indicating that they are not suitable for retrospective assessment of reporting completeness (Rohel et al., 2021; Desmons et al., 2024).

Nonetheless, since the publication of Chipchase et al. (2012), several meta-analytic studies have addressed these limitations by extending the list of items, providing clearer and more concrete definitions for items (Rohel et al., 2021) and/or reporting inter-rater reliability (e.g., Bai et al., 2022; Gunduz et al., 2020; Lanza et al., 2025; Rohel et al., 2021; Snow et al., 2019). However, to date, no unified or updated tool has been developed to integrate these improvements.

### 1.2 Objectives

The overall aim of this study was to build upon Chipchase et al.’s (2012) checklist through an international, community-driven initiative. Specifically, the first aim was to conduct a scoping review to assess current reporting trends, including changes in reporting over time. We examined how frequently the previously identified core TMS methods were reported or controlled, how reliably they could be assessed, and how often the distinction between reported and controlled was used. This ensured that we identified real limitations to address and provided a baseline against which to measure performance. The second aim was to develop a new tool that addresses these limitations. This was done by building on and refining existing items, and generating explicit operational definitions and examples to support consistent interpretation and application. The third aim was to refine and validate the tool on a large body of published TMS studies through multiple phases of development, validation, and test-retest assessment, with revisions informed by qualitative rater feedback and quantitative inter-rater reliability after each phase.

To build, test and validate the tool, we used 333 published studies from the short-latency afferent inhibition (SAI) literature. The SAI protocol involves a relatively complex paired-pulse conditioning design, including electromyography (EMG), peripheral nerve stimulation, and single-pulse TMS, that reflects the methodological demands of a wide variety of TMS studies, was applied to both healthy and clinical populations, and therefore is ideal for this purpose. In addition, we had access to a recently curated systematic review and meta-analysis of this literature, which provided a defined, likely exhaustive coverage of the field (Holmes et al. in preparation), minimising potential bias in the selection of studies.

## 2. Methods

### 2.1 Scoping Review

The study was preregistered on the Open Science Framework (OSF; https://osf.io/tywn8/overview). Searches for the scoping review were conducted in PubMed, Web of Science, and Google Scholar, targeting published studies and grey literature citing Chipchase et al. (2012), and studies that adapted it (Pellegrini et al., 2020; Rohel et al., 2021). Text-based outputs from the databases were combined and processed using custom MATLAB scripts for bibliographic retrieval (*Supplementary Material [SM]1*, https://osf.io/fpu3k/files/rthnv). The last search date was 31 January 2025. Studies from the database searches were included if they were systematic reviews or meta-analyses that provided item-level or study-level reporting based on Chipchase’s checklist.

Screening and data extraction were performed by two independent extractors (NPH and one of sixteen second extractors). Discrepancies between extractors were resolved by NPH, re-checking the extracted data against the original data. Data was extracted using spreadsheet software, including Microsoft Excel, OpenOffice Calc, and Google Sheets. The extracted data included authors, article title, source title (journal or publication), abstract, publication date, volume, issue, part number, start page, end page, article number, the number of studies reviewed, item and study level reporting of Chipchase’s checklist items, and any reported measures of inter-rater agreement. Article level inclusion or exclusion decisions, their reasons, and the database for data extraction, and details of how discrepancies were resolved are available in *SM2* (https://osf.io/fpu3k/files/fqm76). The proportions of studies reporting each checklist item were calculated, and bootstrapped 95% confidence intervals (CIs) were generated (see *SM3* for scoping review analysis code; https://osf.io/fpu3k/files/98e5q).

### 2.2 Tool development

Throughout, we use the term ‘the tool’ to collectively refer to the TMS-RAT, including its reporting items, sections, the guidance document, and the rating spreadsheet. Each version of the tool is numbered, from v0.1 to v0.3 (development), and v1.0 to v1.1 (validation).

#### 2.2.1 Tool design

The first, v0.1 of the tool included all 30 items from the Chipchase checklist, two of which were split into two items (13a: Coil shape, 13b: Coil size; 16a: Coil location, 16b: Coil stability), and two of which were combined (18: Stimulation intensity; 26: Test pulse intensity). Two further items (Cognitive state, Time of day) were taken from Pellegrini et al. (2020), and 49 other items were added by us (e.g., height, experimental design, neuronavigation, threshold, EMG, afferent stimulation, paired pulse, evoked potentials, repetitive TMS). The scoping review informed several design choices, including removing the distinction between ‘reported’ and ‘controlled’ and introducing a distinction between ‘partial’ and ‘full’ reporting to assess reporting more precisely, resulting in three categories (full/part/missing). Whenever available, item definitions were taken or adapted from Rohel et al. (2021), and examples for ‘full’, ‘part’, and ‘missing’ reporting were produced. For quantitative variables such as age or stimulation intensity, ‘full’ reporting was defined as including a measure of both central tendency (e.g., mean or median) and variability (e.g., SD, standard error or interquartile range), to support the quantitative re-use of data in modelling and meta-analysis. We also introduced greater modularity in the tool’s structure and organised items into thematic sections (e.g., participant characteristics, hardware, stimulation intensity) that may be used independently or combined depending on the study’s methods.

The list of items in each tool version and their sources is included in *SM4* (https://osf.io/fpu3k/files/af3pj). Two of the v0.1 sections (paired-pulse—later generalised into the *Conditioning* section—and repetitive TMS; rTMS) were included but not tested during the first development phase. The rTMS section will be validated in a future version. The 9 remaining sections included 67 items (see v0.1 of the tool in *SM5*; https://osf.io/fpu3k/files/5d7bt), and an ∼18,000-word guidance document (*SM6*; https://osf.io/fpu3k/files/e4ucm). Following v0.1, we developed and evaluated v0.2, v0.3 and then v1.0.

#### 2.2.2. Raters

The 17 authors of this paper served as raters during the development, validation, and retest phases. We were at a range of career stages, including PhD students (*n* = 4), postdoctoral researchers (*n* = 8), senior technical specialists (*n* = 1), and more senior academics (*n* = 4); affiliated with 21 different institutions, with only two authors based at the same university and research team. We are based in the UK (*n*=12), the USA (*n*=2), Germany (*n*=1), Canada (*n*=1), and Italy (*n*=1). Seven of us had English as our first language, the language in which the tool was developed and validated. Our individual experience with TMS ranged from 1 to 24 years (*M*=6.5, *SD*=5.5), and the estimated proportion of current research (over the past two years) involving TMS ranged from 15% to 100% (*M*=60.3%, *SD*=30.4%). We represented a wide range of methodological expertise and practical experience, which aligns with the intended and likely user base of the TMS-RAT—a range of researchers actively involved in conducting original studies and systematic reviews.

#### 2.2.3 Tool development phases

All except two raters (OS and NPH, who developed v0.1) were naïve to the tool during their first rating phase. Apart from Phase 1, each further development and validation phase included at least four naïve raters (except phase 3, when one rater withdrew) to assess if the tool could be reliably used by first-time users, and to compare performance with and without prior experience. Raters flagged ambiguities and noted difficulties during rating. This feedback, alongside inter-rater reliability (Gwet’s AC1), informed revisions after each phase. Gwet’s AC1 is a chance-corrected measure of agreement, where a value of 1 indicates perfect agreement, and 0 indicates agreement no better than chance (Gwet 2002, 2014; Vach & Gerke, 2023). Articles were rated in duplicate by pairs of raters matched for experience with previous versions of the tool (naïve vs. not). Within this constraint, rater pairings were rotated across phases to minimise repetition.

Table 1 summarises the number of raters and articles at each phase.

**Table 1.** Number of raters and articles rated across each phase.

| Phase | Version | Raters |  | Articles Rated |  |
| --- | --- | --- | --- | --- | --- |
|  |  | Total | Naïve | Per phase | Cumulative |
| Development 1 | v0.1 | 4 | 2 | 28 | 28 |
| Development 2 | v0.2 | 8 | 4 | 40 | 68 |
| Development 3 | v0.3 | 11 | 3 | 72 | 140 |
| Final Validation | v1.0 | 17 | 6 | 193 | 333 |
| Retest | v1.0/v1.1 | 16 | 0 | 32 | 333 |

Alongside raters’ feedback, we sought open-ended comments from the international expert TMS community. We searched for active TMS researchers in each of 143 countries, querying PubMed for TMS articles published since 1 January 2024, including the affiliated country name. Where possible, one or more co-authors from each paper were identified as active TMS researchers by further examining their PubMed and Google Scholar profiles and their institutional websites. We aimed to contact at least one researcher per country, focusing on non-clinical TMS research. We emailed 86 active TMS researchers from 33 countries to request feedback (*SM7*; https://osf.io/fpu3k/files/rz6nb). Items and sections of the tool were revised based on feedback received.

The tool was distributed and completed by raters using a Google Sheet. From v0.2 onwards, a built-in validation page flagged missed items, prompting raters to complete any omissions. Raters recorded item responses on a three-point scale: 0 for not reported, 0.5 for partially reported, and 1 for fully reported, and provided qualitative feedback comments. Raters were asked to rate each paper in one sitting when possible.

After each phase, Gwet’s AC1 was calculated using the irrCAC package (Gwet, 2019) in R (analysis code provided in *SM8;* https://osf.io/fpu3k/files/pwg68). Mean completion times were estimated using a macro within the rating spreadsheet which recorded the time of each item response; qualitative feedback was reviewed to guide the refinement, combination, or removal of items. After applying pre-registered thresholds for inter-rater reliability (Gwet’s AC1 < 0.60, and/or *p* > 0.05, and/or a lower 95% CI bound < 0.50), we finalised v1.1 for retrospective reporting assessment.

## 3. Results

### 3.1 Scoping review

A total of 513 records citing Chipchase et al. (2012) were retrieved across three databases: PubMed (85), Web of Science (187), and Google Scholar (241). After removing duplicates, 267 unique records remained, 32 of which met the inclusion criteria. These 32 systematic reviews or meta-analyses rated a total of 681 individual TMS studies using the Chipchase checklist. Search logs and MATLAB analysis code are provided in *SM9* (https://osf.io/w9rtz). Figure 1 shows that reporting completeness increased gradually over time, with no clear change following the publication of Chipchase’s (2012) checklist in *Clinical Neurophysiology*. Reporting practices did not vary substantially across journals. Ten journals published ten or more primary research papers. *Clinical Neurophysiology* published the most (47), but reporting was not significantly higher there than in the other nine journals (e.g., vs. lowest reporting *Neurology*, *t*(60) = 1.86, *p* = .084), and there was no evidence of significantly higher reporting in the five years after versus before the checklist (2013-2017, *M*±*SD* = 70.0±12.3%; 2007-2011, *M*±*SD* = 59.9±18.3%, *t*(25) = 1.77, *p* = .089; *SM10*; https://osf.io/fpu3k/files/jwrvh). In addition, while the Chipchase checklist distinguished between items being ‘Reported’ versus ‘Controlled’, only 31% (10/32) of reviews used this distinction.

**Figure 1.**
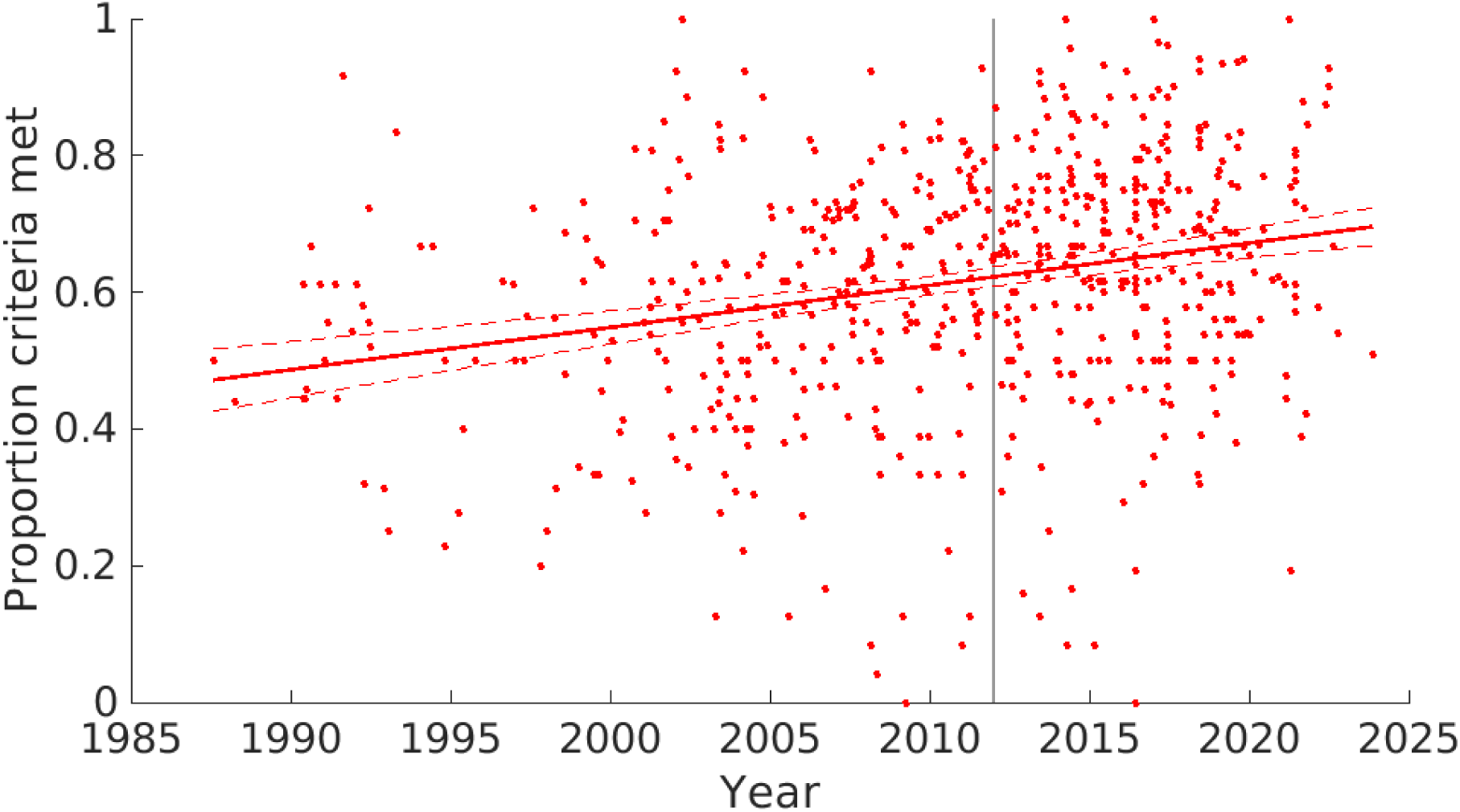
Reporting completeness over time based on Chipchase’s checklist. Individual study reporting and linear trend (with 95% CI) of the proportion of Chipchase items reported over time. Grey line: year of Chipchase et al. (2012) publication. N=614 studies with item-level ratings.

The proportion of studies meeting individual checklist criteria varied substantially (Figure 2). For example, participant age and sex (items #1 and #2), were reported about 90% of the time, while participants’ attention and arousal during testing (item #23), were rarely reported (∼18%). All studies citing Chipchase et al. (2012) and the item-level reporting for all 30 items are provided in *SM11* (https://osf.io/fpu3k/files/jwrvh).

**Figure 2.**
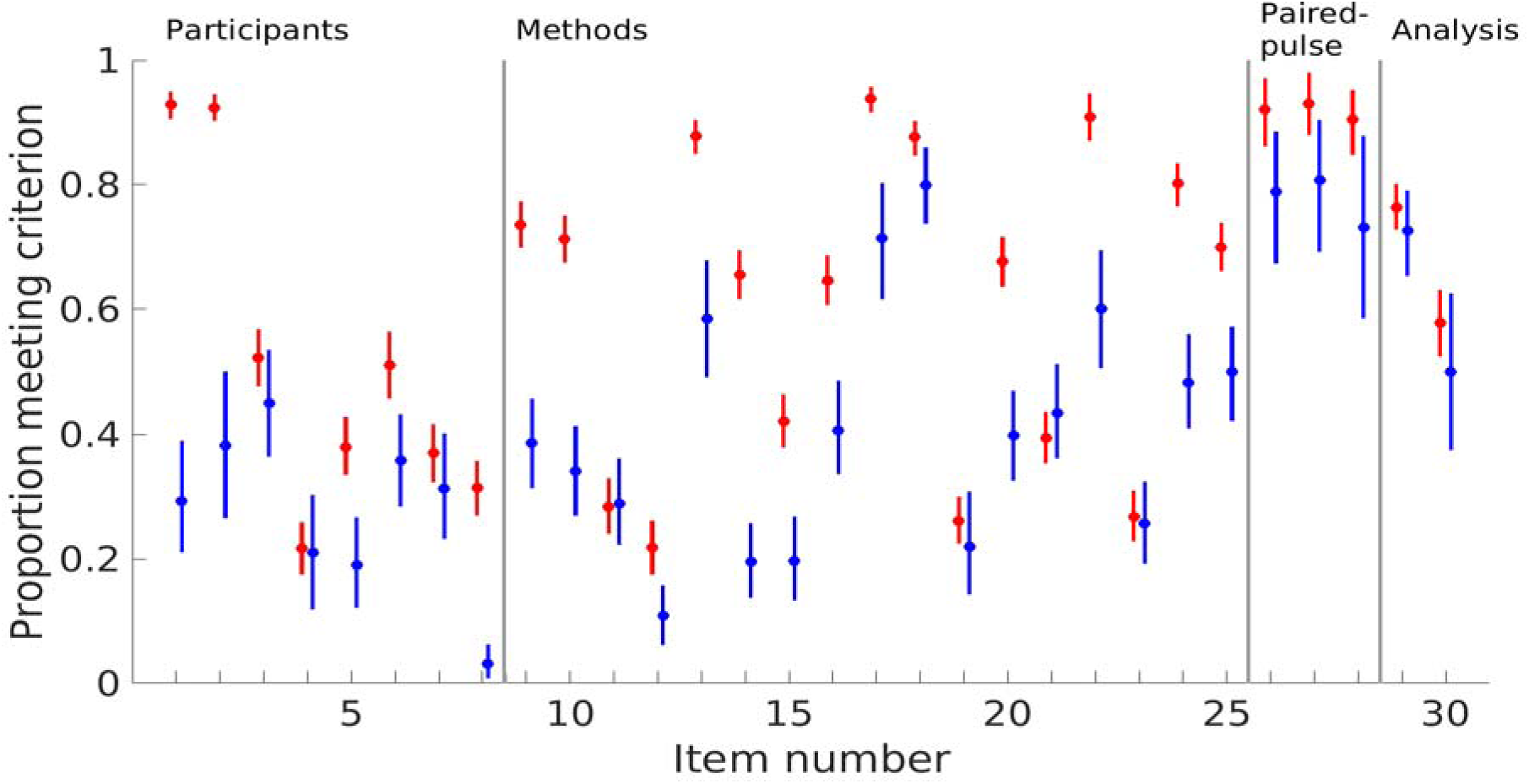
Proportions of 614 studies meeting Chipchase criteria ‘reported’ vs. ‘controlled’. *Note.* Each circle shows the proportion of studies that either reported (red) or controlled for (blue) each item. Items are grouped into thematic sections: Participants, Methods, Paired-pulse, and Analysis. Error bars represent bootstrapped 95% CIs.

### 3.2 Tool development

#### 3.2.1 Feedback from TMS researchers

We received feedback from 12 TMS experts worldwide. The process was open-ended, and contributions varied in scope and depth. Some focused on clarifying item definitions, while others prompted the inclusion and/or removal of items. These contributions played an important role in refining the tool to meet the practical needs of the wider research community. In addition to the feedback that has been used to improve the tool’s relevance and usability, a high proportion of consultations focused on the inclusion of items to assess methodological quality or risk of bias. As the intended scope of the TMS-RAT is exclusively a reporting, rather than quality-assessment, tool, this type of feedback could not be incorporated. Whenever item definitions or examples were changed to reflect feedback, this is noted in the corresponding Supplementary Material.

#### 3.2.2 Inter-rater reliability

Inter-rater reliability increased across development phases (Table 2), while the proportion of fully reported items remained stable at approximately 50% (range: 46–56%). Mean completion time per item decreased slightly across phases, indicating some improvement in usability. In the final validation phase, AC1 did not differ between experienced and naïve raters *t*(13.56) = −1.13, *p* = .28, indicating that the usability was independent of experience with the tool. The number of missing ratings also decreased after introducing the validation page in v0.2.

**Table 2.**
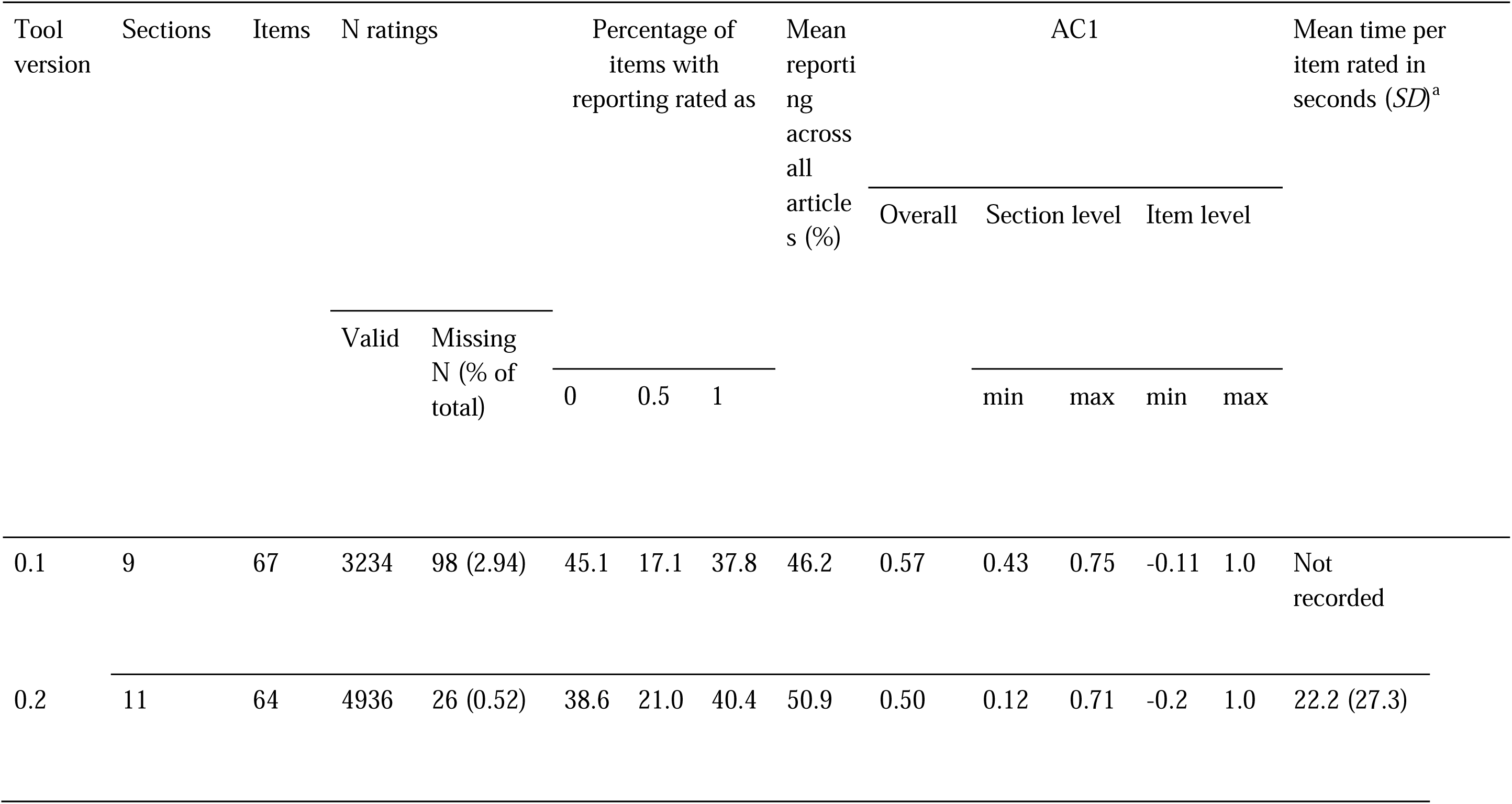
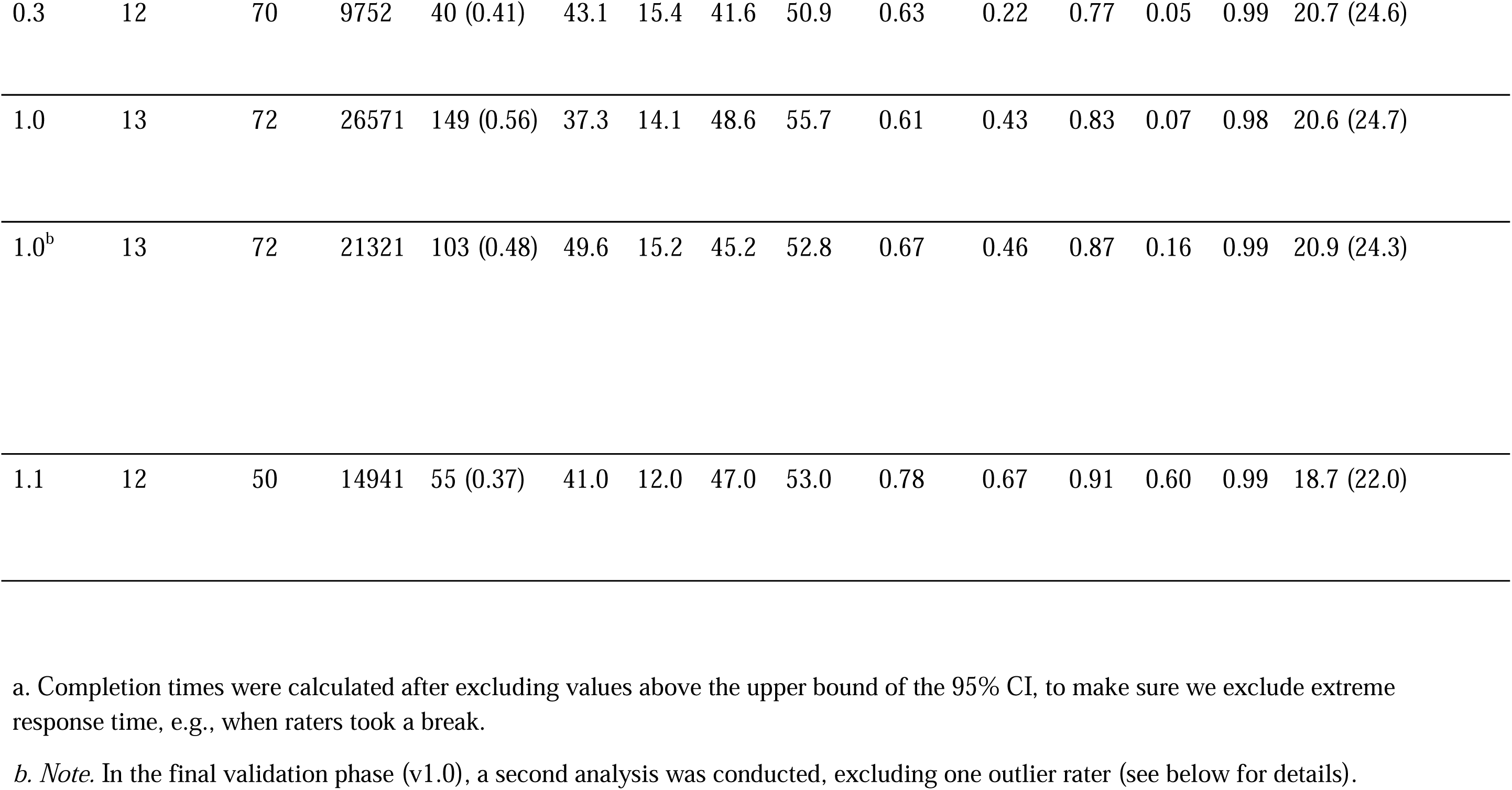
Summary of tool performance across development and validation phases.

In the final validation phase, most raters showed comparable performance (mean AC1 = 0.63, *SD* = 0.08). One outlier showed substantially lower agreement (AC1 = 0.40, *Figure 3*) alongside a high percentage of missing timestamps (53.8%; no other rater had >1%) and markedly shorter completion times (6.93 seconds per item vs. group mean of 20.88, *SD* = 9.36 seconds), indicating that this rater used a different strategy to others. Consequently, this rater was excluded.

**Figure 3.**
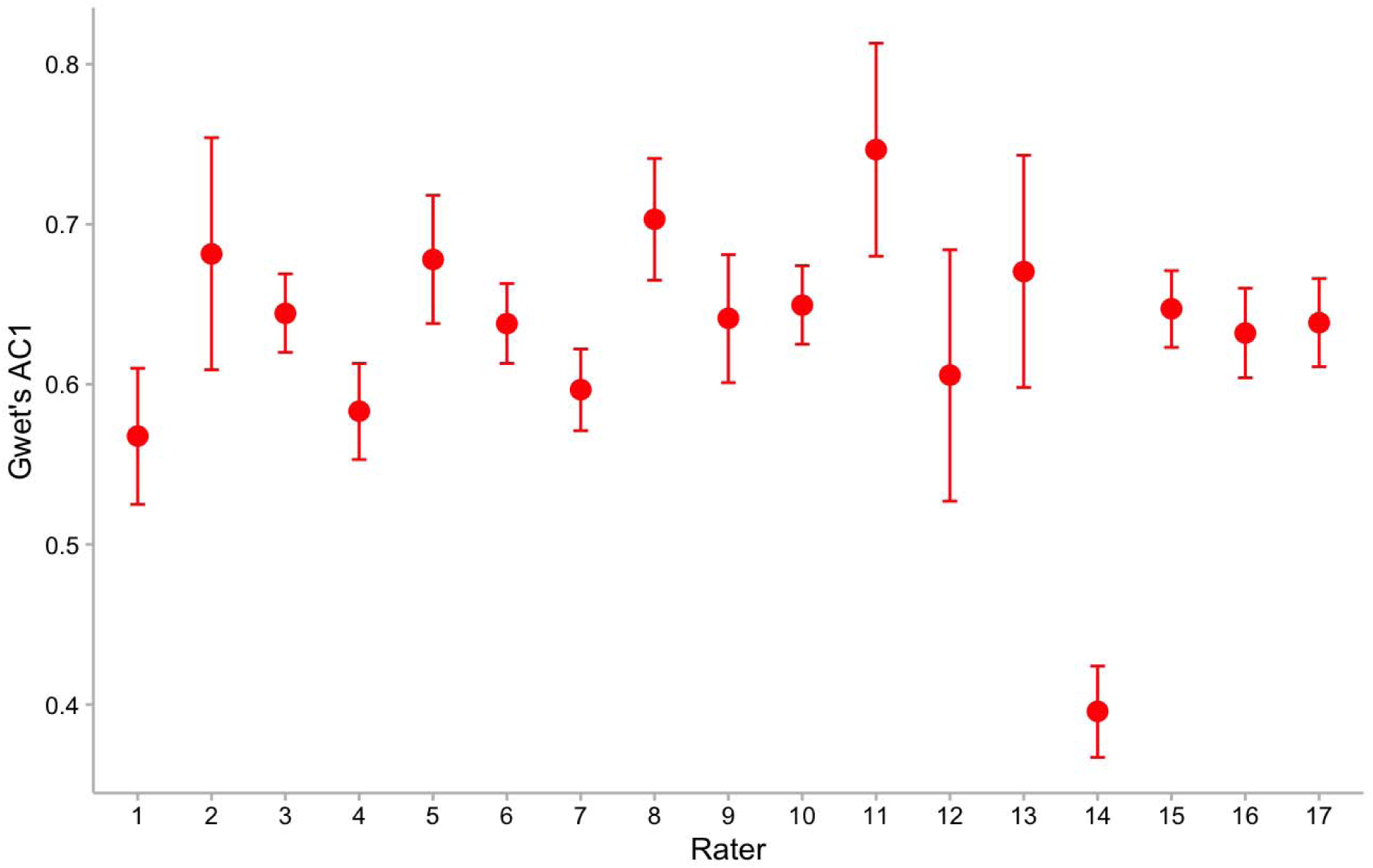
Gwet’s AC1 values for individual raters. Error bars represent 95% CIs.

**Figure 4.**
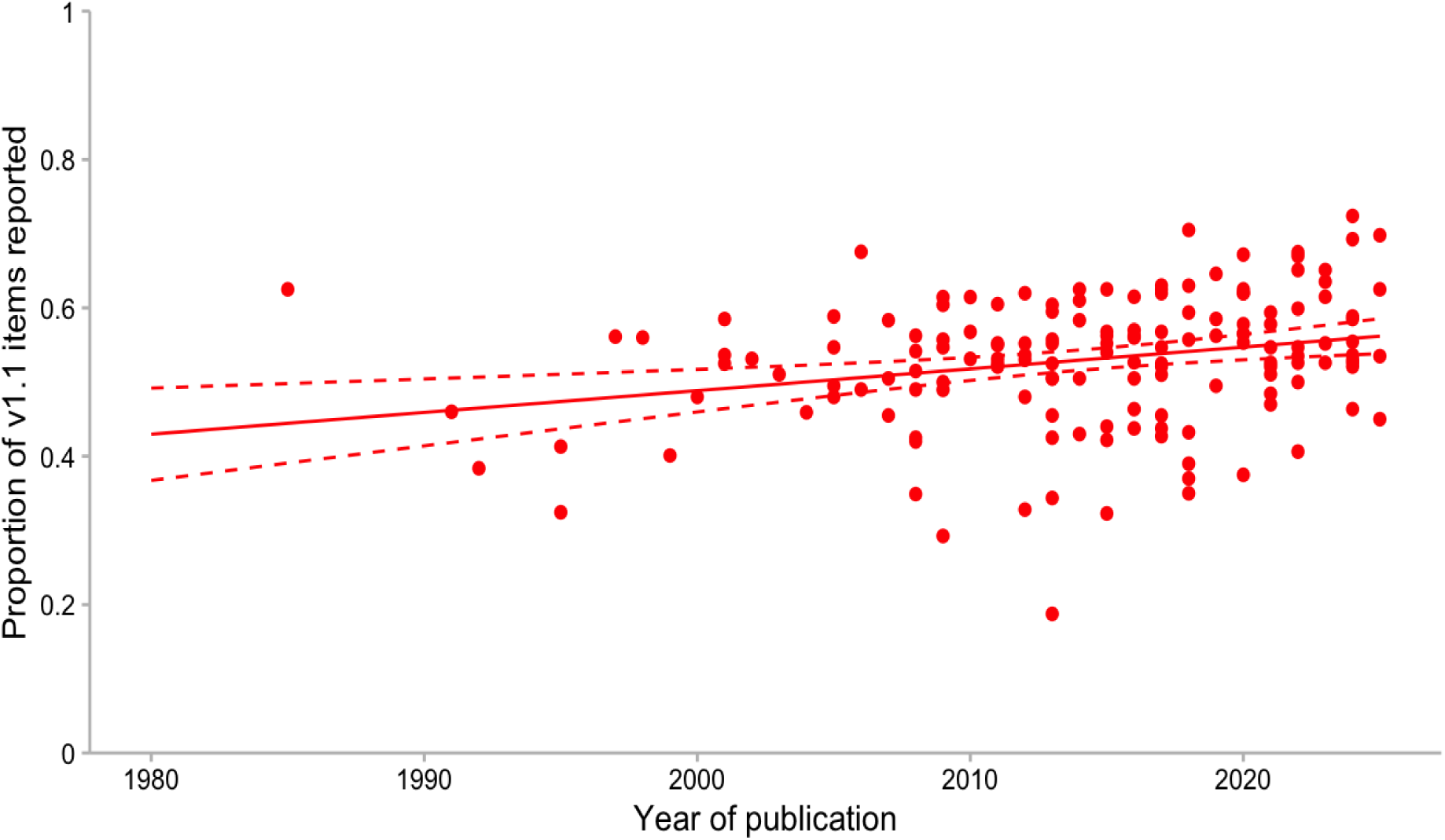
Reporting completeness over time, defined as the proportion of TMS-RAT v1.1 items reported per study. The solid line shows the linear trend, and the dashed lines indicate the 95% CI. *N* = 513 studies with double ratings. For each article, the reported proportion represents the mean proportion of the two raters’ ratings.

#### 3.2.3. Version 1.1: Reliable items for retrospective assessment

The validated version of the tool (v1.0) was filtered using pre-registered criteria (Gwet’s AC1 < 0.60 and/or a *p*-value comparing agreement to chance of *p* > 0.05 and/or a lower 95% CI bound < 0.50). Twenty-two items did not meet these criteria (see *SM12* [https://osf.io/fpu3k/files/vnw56] for the full list and *SM13*[https://osf.io/q3zck] for discussion of why agreement could not be reached), indicating that it is not currently possible to provide a single version of the TMS-RAT that both guides reporting of all key methods and includes only items that can be reliably assessed retrospectively. The final version of the tool, excluding these 22 items (v1.1), is appropriate for retrospective reporting assessment. It includes 12 sections and 50 items (excluding the unvalidated rTMS section). See Table 3 for the full list of v1.0 and v1.1 items.

**Table 3.**
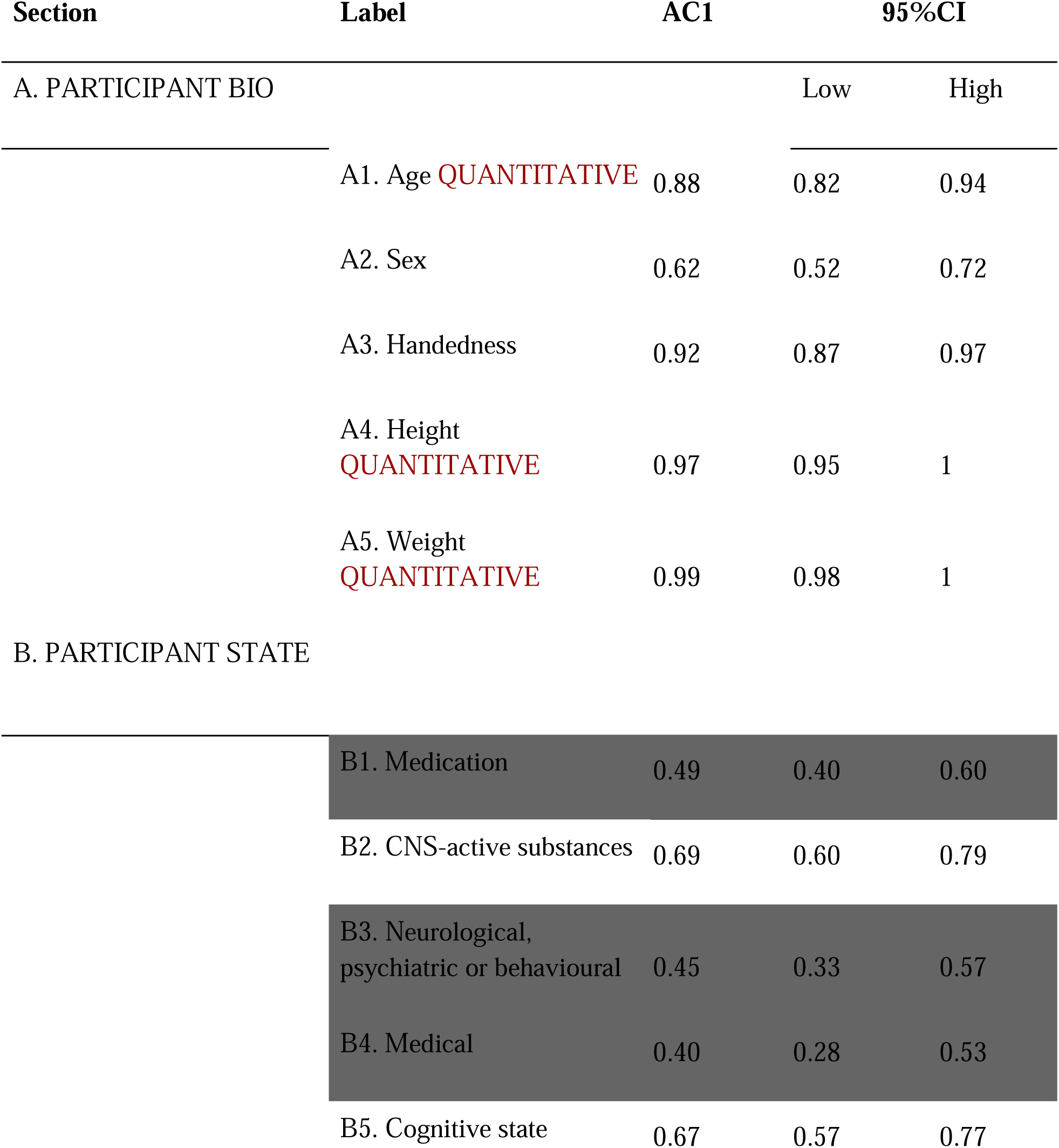
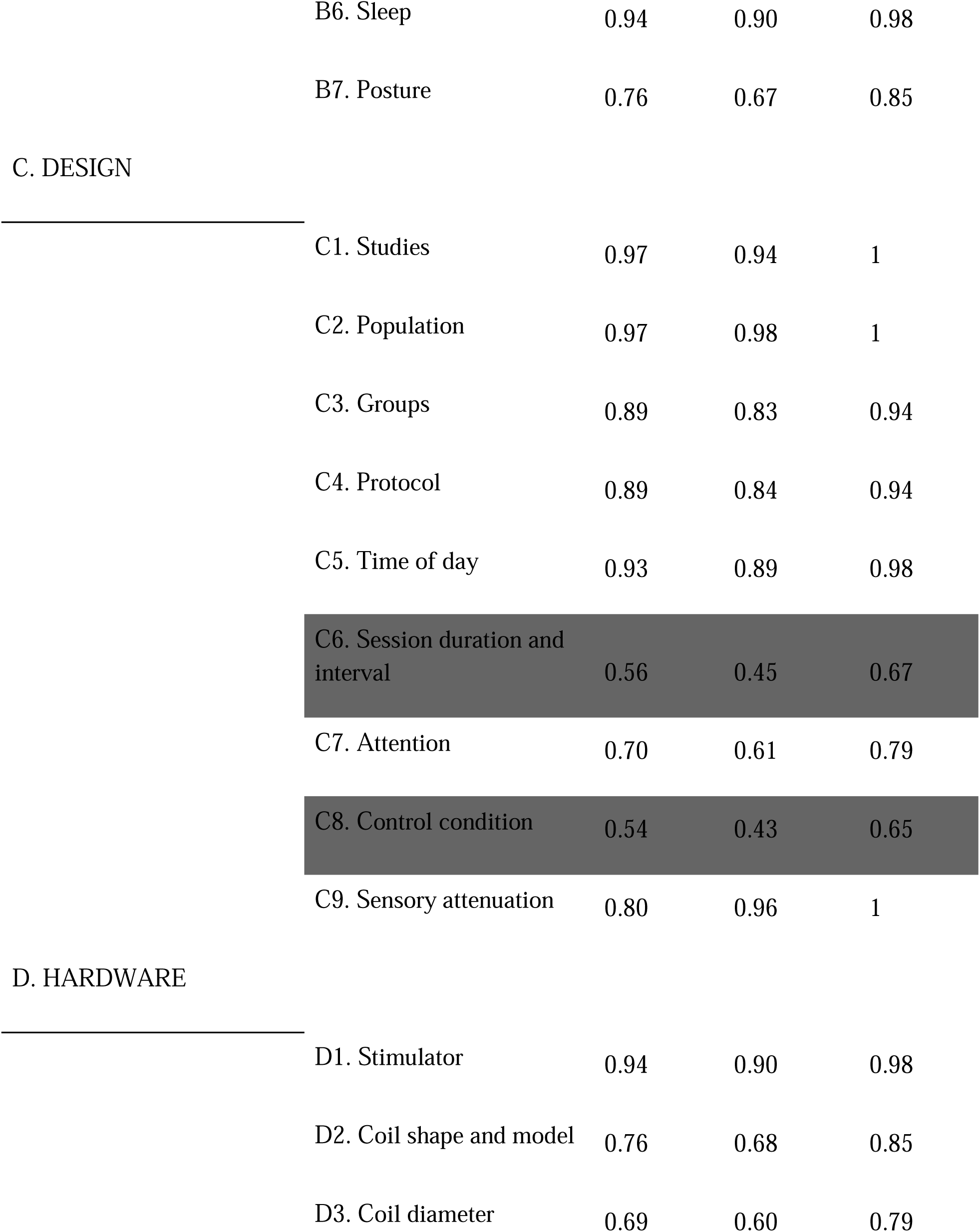
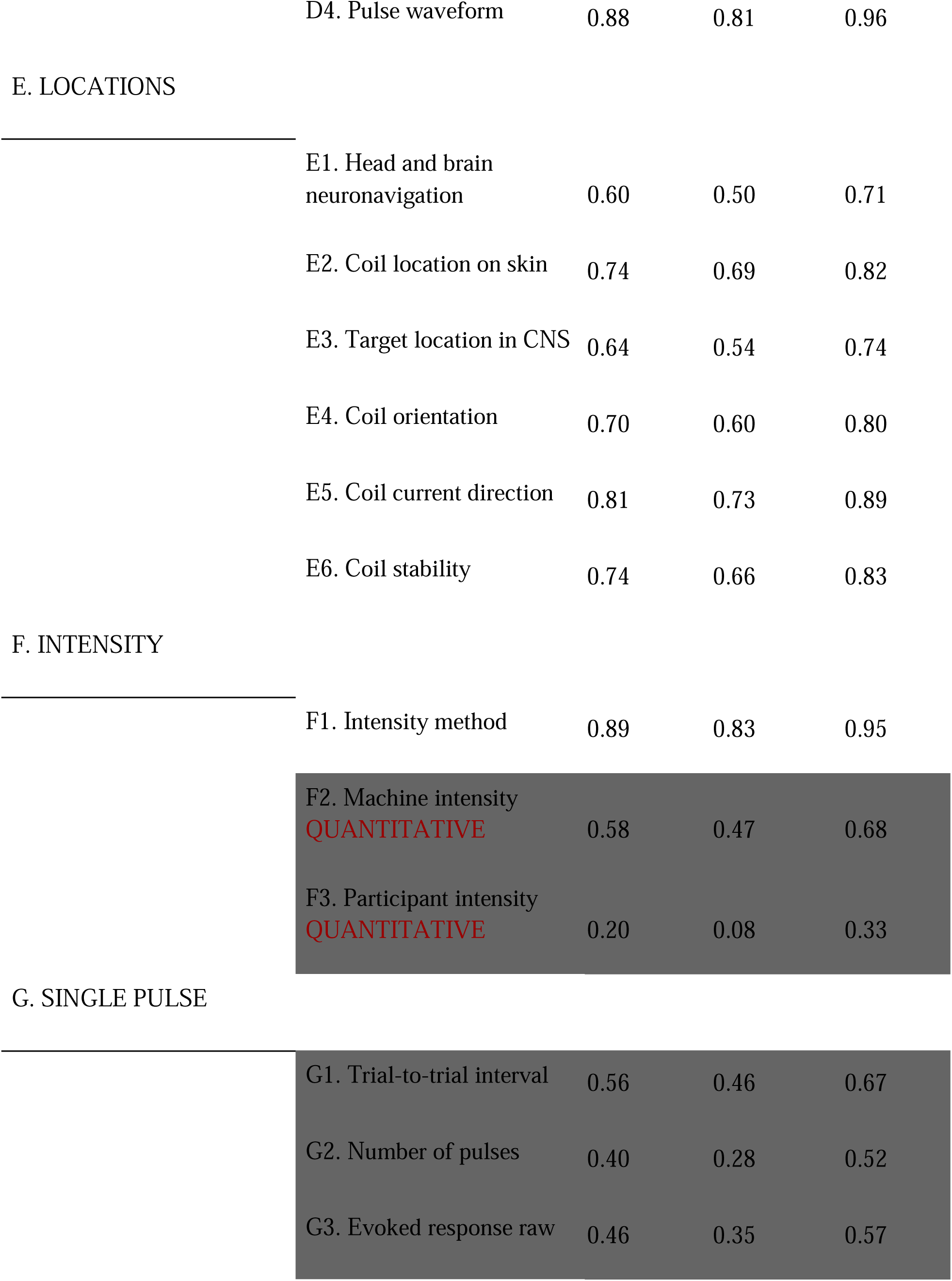
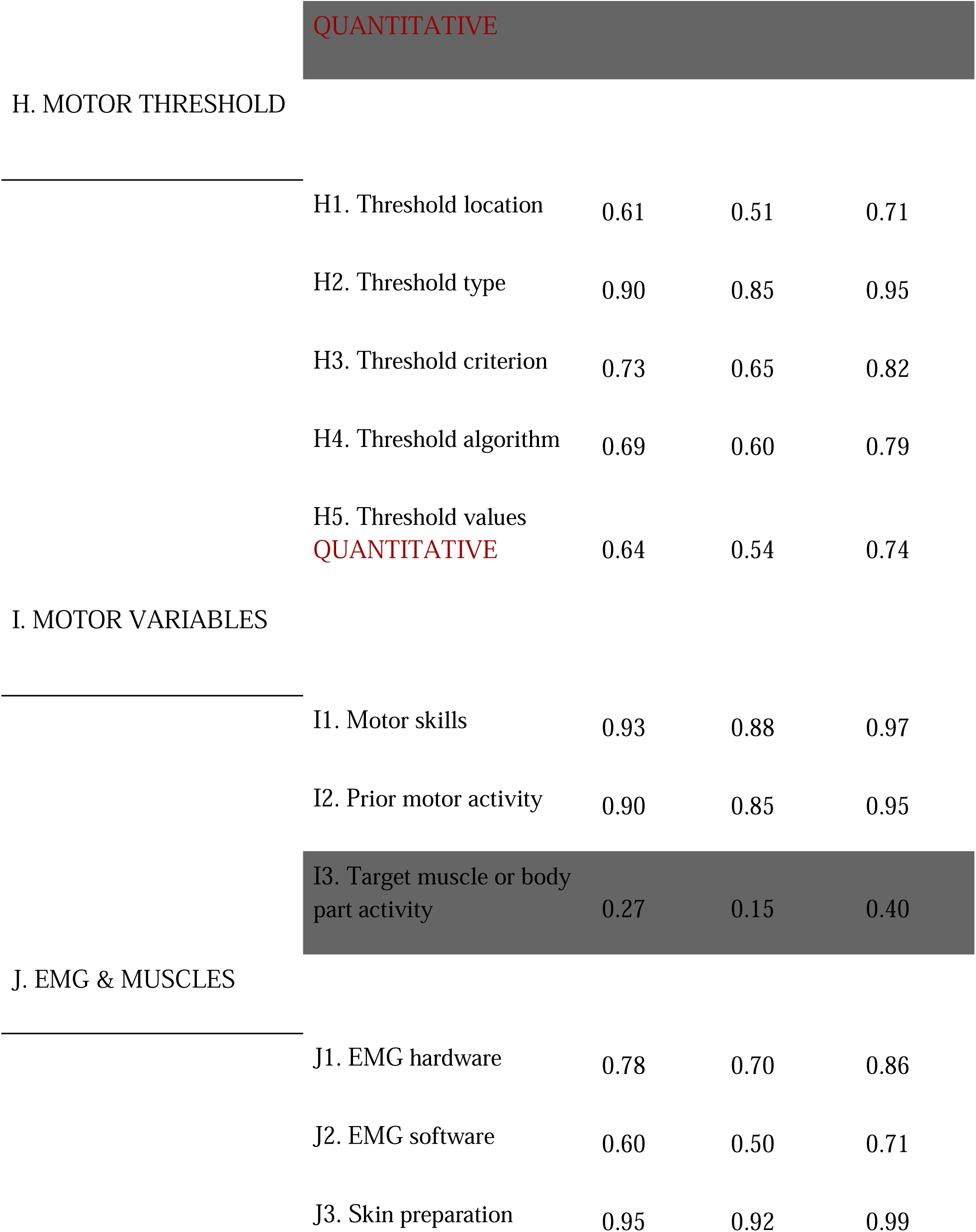
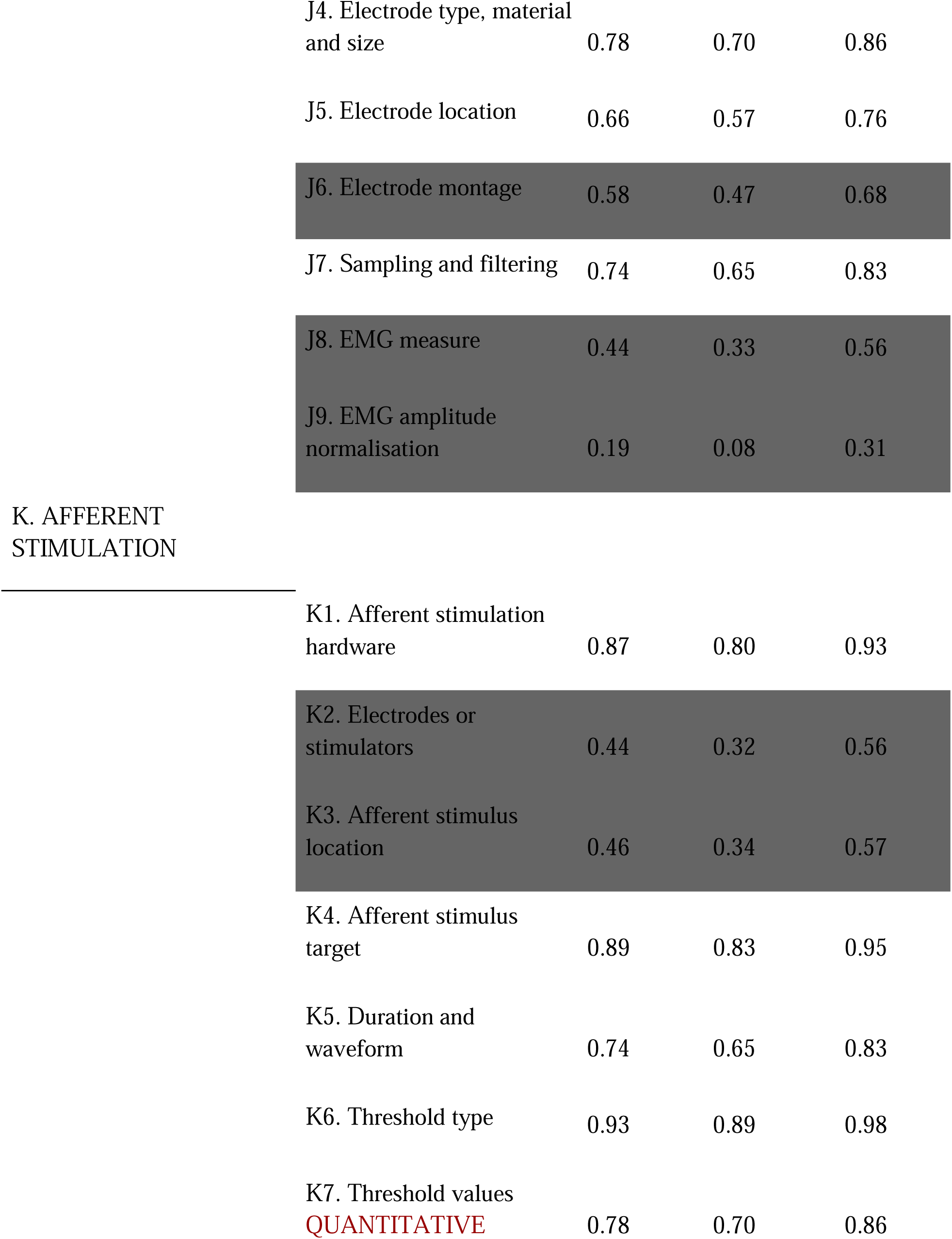
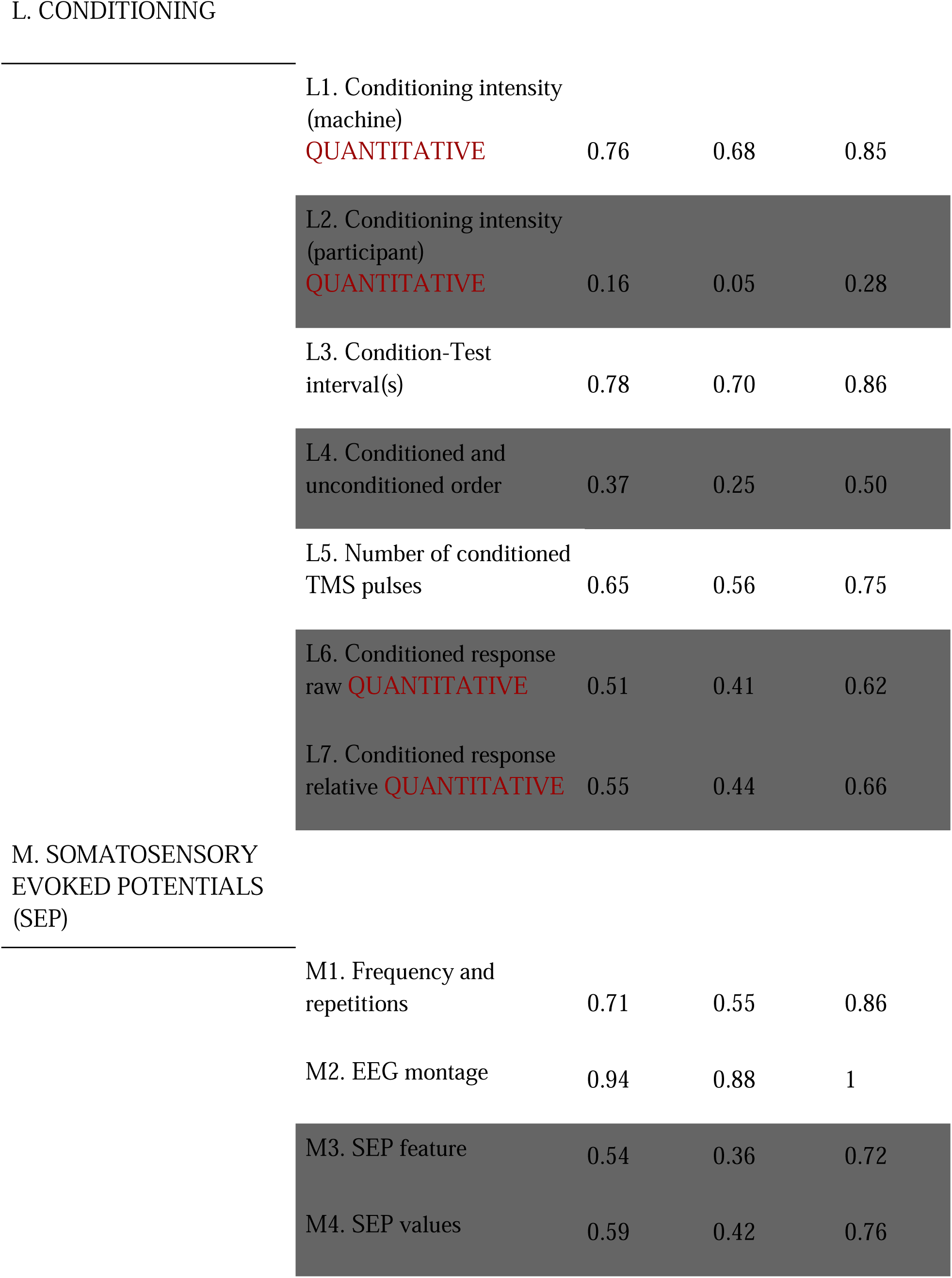
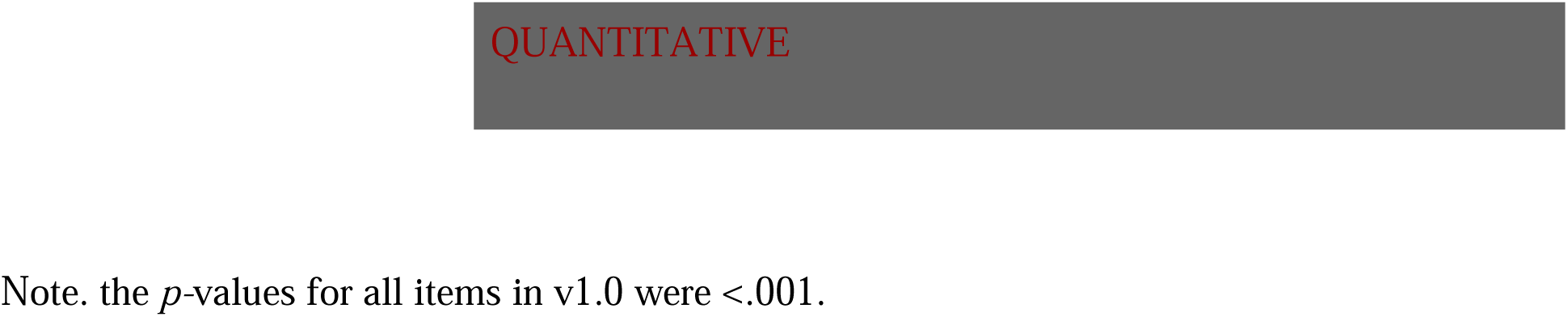
Full list of v1.0 items. Items excluded from v1.1 are shown in dark grey.

#### 3.2.4 Overall reporting trends

Using v1.1 of the tool, we explored overall trends of reporting in the literature. Over time (by publication date), the mean rating per paper showed a gradual improvement β = 0.29%, 95% CI: 0.12 to 0.47%, *p* = .0013. This resembles the, even stronger, positive trend observed for the Chipchase checklist (Figure 1) over time, β =0.62%, 95% CI = 0.43-0.80%, *p* <.001. However, in the TMS-RAT v1.1, the distribution was far from floor or ceiling effects (range 0.19 to 0.72), as 0% of papers had <15% or >85% of items reported, unlike the distribution for Chipchase’s checklist, where 9% of papers had <15% or >85% of items reported.

#### 3.2.5 Test-retest reliability

Test-retest reliability was assessed by re-rating four articles per rater (64 in total) using version 1.0 of the tool, with a minimum interval of two months between the original test and the retest. Analyses were restricted to the subset of items retained in TMS-RAT v1.1. Overall test-retest reliability of the tool was Gwet’s AC1 0.82 (95% CI: 0.80 to 0.84). Item-level AC1 values ranged from 0.65 to 1.00, while section-level AC1 values ranged from 0.68 to 0.96.

#### 3.2.6 Documentation of each development and validation phase

All data and documentation for each phase of tool development and validation are available in the SMs. For v0.1, this includes the raw rating data (*SM14*; https://osf.io/fpu3k/files/yfjca), item-level inter-rater reliability (*SM15*; https://osf.io/fpu3k/files/3m8eh; timing not recorded), and section-level inter-rater reliability statistics (*SM16*; https://osf.io/fpu3k/files/k4jmp). For v0.2, this includes the tool (*SM17*; https://osf.io/fpu3k/files/nxvka), guidance document (*SM18*; https://osf.io/fpu3k/files/zkhdq), a summary of changes since v0.1 (*SM19;* https://osf.io/b6ftz), raw rating data (*SM20*; https://osf.io/fpu3k/files/mzuy6), item-level inter-rater reliability and timing results (*SM21*; https://osf.io/fpu3k/files/ysvch), and section-level inter-rater reliability results (*SM2*2; https://osf.io/fpu3k/files/vx4cm). For v0.3, this includes the tool (*SM23*; https://osf.io/fpu3k/files/z4tqf), the guidance document (*SM24*; https://osf.io/fpu3k/files/9w58f), the summary of changes from v0.2 (*SM25;* https://osf.io/fpu3k/files/nfqxs), the raw rating data (*SM26*; https://osf.io/fpu3k/files/jcpqg), the item-level inter-rater reliability and timing results (*SM27*; https://osf.io/fpu3k/files/bhmn5), and the section-level inter-rater reliability results (*SM28*; https://osf.io/fpu3k/files/n3qcp). For the final validation version (v1.0), we also provide the tool (*SM29*; https://osf.io/fpu3k/files/uvby9), the guidance document (*SM30*; https://osf.io/fpu3k/files/ga9xc), the summary of changes (*SM31;* https://osf.io/b6ftz), the raw rating data (*SM32*; https://osf.io/fpu3k/files/e8ms7), the item-level inter-rater reliability and timing results (*SM33*; https://osf.io/fpu3k/files/26zwe), and the section-level inter-rater reliability results (*SM34*; https://osf.io/fpu3k/files/ypzsf). Following exclusion of the outlier rater, updated item- and section-level inter-rater reliability results for v1.0 are provided in *SM35* (https://osf.io/fpu3k/files/hsd6z) and *SM36* (https://osf.io/fpu3k/files/xb3p9), respectively. For the final, retrospective version (v1.1), we provide the tool (*SM37;* https://osf.io/fpu3k/files/akgwd), item-level inter-rater reliability and timing results (*SM38*; https://osf.io/fpu3k/files/bxrpq), and section-level inter-rater reliability results (*SM39*; https://osf.io/fpu3k/files/ugnjm). The test-retest phase raw rating data is available in *SM40*(https://osf.io/fpu3k/files/6cm5f) (reliability AC1 values are available in *SM41* (https://osf.io/fpu3k/files/spzjc), while section-level AC1 values in *SM42 (*https://osf.io/fpu3k/files/9pfaj).

## 4. Discussion

### 4.1 Summary of Contribution

After assessing current reporting trends in TMS research, we developed and validated the TMS-RAT through iterative testing and refinement, across 333 papers rated by 17 independent raters, and consultation with 12 TMS experts worldwide. We established a comprehensive, empirically tested, community-informed tool ready for implementation by TMS researchers. Although the TMS-RAT was developed specifically for TMS studies, it aligns conceptually with broader reporting frameworks such as CONSORT for clinical trials (Hopewell et al., 2025), and PRISMA for systematic reviews (PRISMA-P Group et al., 2015). TMS-RAT v1.0 and v1.1 together provide a framework for improving reporting and the assessment of reporting across a wide range of study designs, and thereby increasing transparency and reproducibility in TMS studies.

The TMS-RAT builds on the Chipchase et al. (2012) checklist by retaining its items selected by Delphi consensus, while addressing its limitations through operational definitions, examples, and empirical evaluation of usability, inter-rater reliability, and test-retest reliability. We introduced a modular structure that allows the tool to be adapted to different research contexts and provides flexibility for extension into further contexts. We removed the ‘reported’ versus ‘controlled’ distinction because the limited uptake suggested that it is not practical or meaningful to researchers. In contrast, distinguishing between full, partial, and missing information allowed reporting to be characterised with greater resolution, avoided floor and ceiling effects, and better reflected the continuum of reporting completeness.

We assessed inter-rater reliability with Gwet’s AC1, which is calculated treating ratings as categorical rather than ordinal, to avoid inflating agreement estimates and to maintain comparability with the categorical scoring used in the Chipchase (2012) checklist. This is a conservative comparison, as the TMS-RAT applies a three-level categorisation compared to the Chipchase et al. (2012) binary categories. However, all items retained in TMS-RAT v1.1 met predefined reliability criteria, whereas approximately one third of Chipchase items did not meet the same criteria (Rohel et al., 2021; Desmons et al., 2024). The similarity in reporting trends over time and the overall reporting completeness suggests that TMS-RAT v1.1 reflects the same underlying reporting practices as the Chipchase checklist.

### 4.2 Constraints on Reporting Assessment

Twenty-two reporting items from v1.0 did not meet our pre-registered reliability threshold and were excluded from v1.1. This included items considered essential, for example, the entire single-pulse section, as well as items such as the EMG electrode montage and the session duration and interval. Despite iterative attempts to produce clearer instructions or examples, raters could not consistently agree on whether the information was reported. This likely reflects inconsistent reporting of protocols in the literature. Until there is broader alignment in how key methods are reported, a tool cannot be both maximally comprehensive and uniformly reliable, and separate tools for prospective and retrospective use are necessary.

### 4.3 The Intended Use of the TMS-RAT

We recommend using TMS-RAT v1.0 as guidance for reporting methods in original research. Authors are encouraged to complete all relevant sections and to include the completed sheets and the output .csv file as supplementary material when submitting manuscripts for peer review. Reviewers and journal editors are encouraged to assess the completeness of study reporting using the tool as a benchmark.

For retrospective assessments, such as in systematic reviews or meta-analyses, we recommend using TMS-RAT v1.1, which contains only items with high inter-rater and test-retest reliability. This version is intended for evaluating and comparing reporting completeness across studies. Overall reporting completeness can be calculated as the proportion of reported items within the study. This metric could be used as a covariate in meta-regression or subgroup analyses. Importantly, neither v1.0 nor v1.1 of the tool is designed to measure methodological quality or risk of bias, but rather to assess the completeness of reporting, and v1.1 summary measures should be interpreted accordingly.

To make the use of the TMS-RAT easy and intuitive, researchers will be able to use our website, (tms-rat.org) to generate a near-complete Methods section in accordance with v1.0; and to retrospectively assess reporting completeness using v1.1.

### 4.4 Future Development

The TMS-RAT is intended as a living resource that will evolve in response to changes in research practice and community feedback. The current version focuses on studies using TMS alone (or in combination with electromyography and somatosensory evoked potentials). Future development will include updates and extensions to currently unvalidated paradigms, including rTMS, paired-associative protocols, clinical applications, and multimodal approaches such as TMS coregistered with functional magnetic resonance imaging and electroencephalography. The latest validated versions of the tool, along with the guidance document, are openly accessible via the project website (tms-rat.org) as well as our GitHub pages (https://github.com/TMSMultiLab/TMSMultiLab/wiki/TMS-Reporting-Assessment-Tool). We encourage the community to contribute to the ongoing refinement of the TMS-RAT by using it both to prospectively generate Methods sections and to retrospectively assess reporting completeness. Real-world usage feedback and data will be incorporated into future versions of the tool.

## Acknowledgement

We thank all researchers who have provided feedback and suggestions for the TMS-RAT (acknowledged on our website).

